# Rapid enzymatic assay for antiretroviral drug monitoring using CRISPR-Cas12a enabled readout

**DOI:** 10.1101/2024.11.25.625292

**Authors:** Maya A. Singh, Megan M. Chang, Qin Wang, Catherine Rodgers, Barry R. Lutz, Ayokunle O. Olanrewaju

## Abstract

Maintaining efficacy of human immunodeficiency virus (HIV) medications is challenging among children because of dosing difficulties, the limited number of approved drugs, and low rates of medication adherence. Drug level feedback (DLF) can support dose optimization and timely interventions to prevent treatment failure, but current tests are heavily instrumented and centralized. We developed the REverse-transcriptase ACTivity-crispR (REACTR) assay for rapid measurement of HIV drugs based on the extent of DNA synthesis by HIV reverse transcriptase. CRISPR-Cas enzymes bind to synthesized DNA, triggering collateral cleavage of quenched reporters and generating fluorescence. We measured azidothymidine triphosphate (AZT-TP), a key drug in pediatric HIV treatment, and investigated the impact of assay time and DNA template length on REACTR’s sensitivity. REACTR selectively measured clinically relevant AZT-TP concentrations in the presence of genomic DNA and peripheral blood mononuclear cell lysate. REACTR has the potential to enable rapid point-of-care HIV DLF to improve pediatric HIV care.

## Introduction

Over 40% of the estimated 1.4 million children living with HIV worldwide had access to antiretroviral therapy (ART) in 2023.^1^ Even among those with access, pediatric ART dosing is challenging to implement – often done by weight and requiring multiple pills instead of fixed dose combinations – and children’s rapid growth and developmental changes affect drug metabolism and pharmacokinetics.

Moreover, given the added safety constraints associated with implementing clinical trials among children, there are fewer approved ART regimens for children.^2–4^ The scarce amount of safety and pharmacokinetic data for children and neonates slows the approval of new treatments, which increases the risk of drug resistance.^2,5,6^ There is a growing need for age-appropriate formulations and precise dosing to ensure therapeutic efficacy and minimize adverse effects.^7^

In addition, rates of non-adherence to medication among children are very high, compounding the difficulty of pediatric dosing. Non-adherence arises due to factors such as complex medication regimens, side effects, inadequate support systems, and socioeconomic barriers.^8^ To ensure consistent and proper medication intake in children, many tailored strategies have been developed and piloted, such as caregiver support, simplified treatment regimens, and interventions addressing social and behavioral factors, but have been limited in the success of improving adherence.^9^ The efficacy of these interventions is often assessed with adherence monitoring strategies like surveys and pill counts, but these subjective measures do not always correlate with treatment outcomes. In contrast, quantitative HIV viral load monitoring is a lagging index of non-adherence and may not indicate impending failure until symptoms and drug resistance have occurred.

HIV drug level feedback (DLF) provides objective information about medication adherence that correlates with treatment outcomes. Clinical trials, primarily among adults, have established drug levels that correspond to different adherence thresholds and treatment outcomes like virological failure. DLF has also been useful among other key populations such as adolescent girls and young women (AGYW), where both AGYW and their healthcare providers express a desire for routine DLF with concomitant increases in adherence and HIV prevention outcomes.^10–12^

Despite the promise of HIV DLF, its use has largely been restricted to clinical and implementation trial settings because of the high cost and heavy instrumentation required to perform gold standard liquid chromatography tandem mass spectrometry (LC-MS/MS) measurements.^13–15^ Although LC-MS/MS results were provided to AGYW and their healthcare providers in studies like MTN-034/REACH, the typical time to results was ∼56 days, significantly delaying interventions.^10,12^ Recently developed point-of-care (POC) tests, like the urine tenofovir (TFV) lateral flow assay (LFA) provide a rapid and inexpensive alternative and also increased adherence to pre-exposure prophylaxis (PrEP) regimens among AGYW in Kenya.^11,16,17^ However, tenofovir-based regimens are not approved for use in children and LFAs have not yet been developed for the key antiretroviral drugs used in pediatric regimens, namely azidothymine (AZT), abacavir (ABC), and lamivudine (3TC).^9^ Thus, there is a need for new tests for HIV DLF among children in POC settings. Such tests would find immediate use in monitoring and improving adherence and precision/personalization in pediatric ART dosing to increase efficacy and reduce toxicity.

We recently developed REverse transcriptase ACTivity (REACT) assays for rapid and minimally instrumented measurement of reverse transcriptase (RT) inhibitors – the backbone of HIV treatment and prevention regimens – as a function of the drugs’ activity.^18–21^ We have shown that REACT can measure clinically relevant concentrations of the active intracellular forms of RT inhibitors including TFV diphosphate (TFV-DP), 3TC triphosphate (3TC-TP), and AZT triphosphate (AZT-TP).^18–20^ We have also demonstrated that by designing custom DNA templates based on the chemical structure of nucleotide analog drugs and Watson-Crick-Franklin base pairing, we can measure the activity of co-administered drugs without cross-reactivity.^20^ We expanded the capabilities of this assay to measure multiple non-nucleotide analog RT inhibitors with readout in a portable and inexpensive ($300) fluorescent reader.^21^

Despite the potential of REACT to provide a path to POC HIV DLF, there are key technical challenges that prevent its direct application to pediatric ART monitoring. Our prior implementation of the assay relied on an intercalating dye to provide non-specific fluorescence when bound to double-stranded DNA (dsDNA) synthesized by HIV RT. For drugs like TFV-DP that accumulate in RBCs, REACT can measure DNA synthesis with minimal interference from background genomic DNA (gDNA) since RBCs do not have a nucleus. However, all the approved antiretroviral drugs for children do not accumulate appreciably in RBCs and must be measured in peripheral blood mononuclear cells (PBMCs), where background gDNA would interfere with the assay signal.^9,22,23^ Furthermore, the intercalating dye non-specifically inhibited HIV RT activity and required endpoint addition, which introduced an additional timed assay step while preventing the acquisition of real-time assay information that could prove vital in assay design and optimization.

Here we present REACT with CRISPR-enabled readout (REACTR) assays where the non-specific intercalating dye readout is replaced by CRISPR Cas complexes that target the sequence of synthesized DNA generated by HIV RT. REACTR leverages advances in CRISPR-based diagnostics that enable high sensitivity and specificity, versatility, and ease of use in measuring nucleic acid targets.^24–28^ Specifically, CRISPR-Cas12a is a powerful gene-editing tool that uses a Cas12a enzyme and a guide RNA (crRNA) to precisely target DNA at specific locations with attomolar sensitivity.^29^

Additionally, Cas12a can cut nearby DNA after activation through its collateral cleavage activity, thus enabling fluorescent readout.^30^ We demonstrate that REACTR can provide real-time information about HIV RT activity in the presence of AZT-TP and characterize the impact of design parameters like reaction time and DNA template length on REACTR’s sensitivity. We optimize REACTR to measure AZT-TP concentrations that are clinically relevant for ART adherence and dosing in children and validate assay performance in more complex sample matrices, including gDNA and PBMC lysate, where our previous intercalating dye readout failed.

## Results & Discussion

### REACTR Overview and Process

REACTR measures DNA synthesis by HIV RT using a CRISPR-Cas12a reporter system. We engineered a DNA template such that the complementary DNA (cDNA) synthesized by HIV RT activates the crRNA-Cas12a complex and generates fluorescence through collateral cleavage of a reporter molecule. Nucleotide RT inhibitors (NRTIs) and deoxynucleotide triphosphates (dNTPs) compete for incorporation into cDNA (Figure 1A). At high NRTI concentrations, cDNA chain termination is more favorable, resulting in short DNA fragments that do not contain the crRNA-Cas12a recognition sequence, leaving the Cas12a complexes in the reaction inactivated. Conversely, at low NRTI concentrations, full-length cDNA synthesis is more likely, resulting in long DNA fragments with intact crRNA-Cas12a recognition sequences and high levels of Cas12a complex activation. Thus, the amount of Cas12a activation, as measured through fluorescence, can be used to infer NRTI concentrations (Figure 1B).

**Figure 1.**
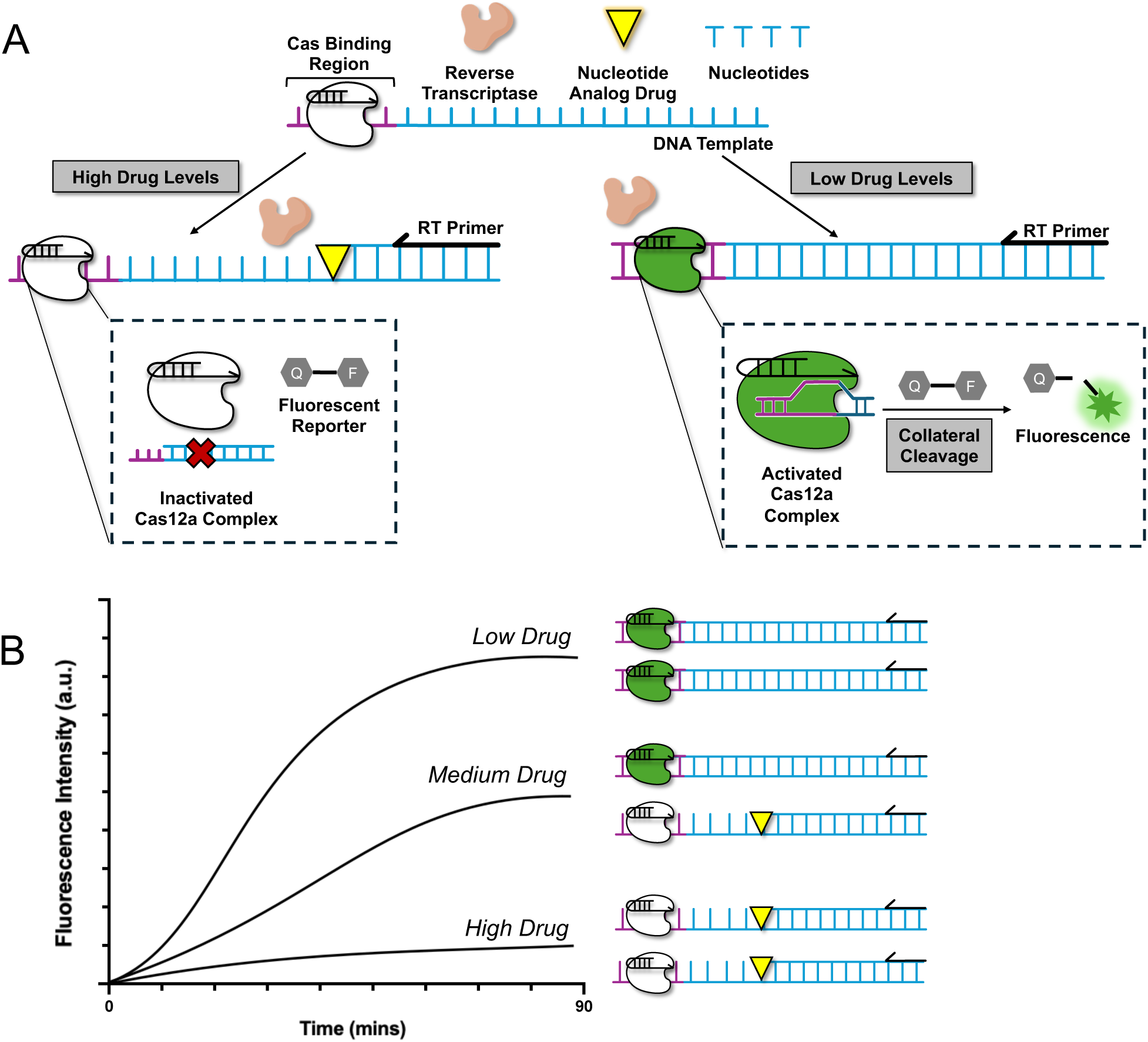
REACTR Assay Overview. **(A)** The REACTR assay measures the concentration of nucleotide reverse transcriptase inhibitors (NRTIs) as a function of DNA synthesis by HIV reverse transcriptase (RT). At high NRTI levels, DNA chain termination occurs and inactivated Cas12a complexes are unable to generate fluorescence. However, at low NRTI levels, Cas12a complexes are activated and cleave reporter complexes leading to increased fluorescence. **(B)** An example schematic of real-time fluorescence curves obtained with REACTR at low, medium, and high NRTI concentrations.

### REACTR accurately measures AZT-TP concentrations in buffer

We tested the REACTR assay’s ability to measure AZT-TP, the active intracellular metabolite of zidovudine – the first ever antiretroviral drug used for HIV treatment and prevention, which is still in use in pediatric treatment regimens. We spiked AZT-TP into aqueous buffer at concentrations (1E-4 M to 1E-10 M) spanning 3 orders of magnitude above and below the physiologically relevant range and added those samples into REACTR assays with real-time fluorescence measurement for 90 minutes (Figure 2A). For reactions containing AZT-TP concentrations >1E-7 M, DNA synthesis was inhibited such that fluorescent curves were comparable to the negative controls (no RT, no drug). At AZT-TP concentrations ≤1E-7 M, fluorescence intensity increased over time, indicating that DNA synthesis by HIV RT activated Cas12a complexes. As expected, higher fluorescence values were obtained at lower AZT-TP concentrations with the lowest drug concentration tested (1E-10 M) overlapping with the positive (RT, no drug) control.

**Figure 2.**
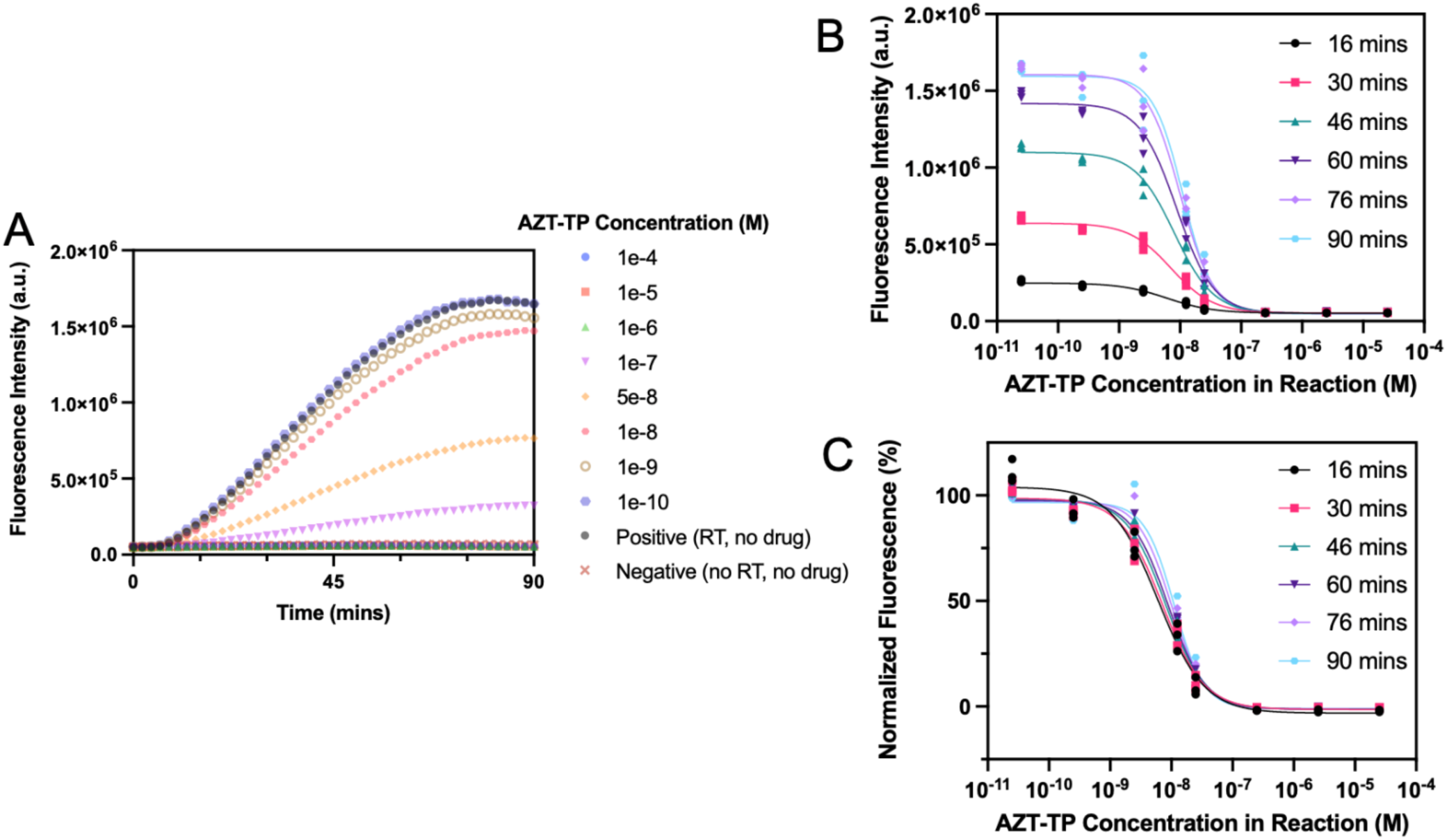
REACTR Real-time Performance and Characterization with AZT-TP. **(A)** Real-time REACTR curves with varying concentrations of AZT-TP. Fluorescence measurements were collected every 2 minutes. Each data point represents the mean fluorescence measurement of three replicates. Supplementary Figure 1 shows this data set in triplicate. We generated inhibition curves from **(B)** raw fluorescence and **(C)** normalized fluorescence as a function of AZT-TP concentrations at different time points (N = 3).

We created classic sigmoidal enzyme inhibition curves for data at ∼15-minute increments (Figure 2B). The inhibition curves with data points at 76 and 90 minutes overlapped curves because the reaction had plateaued. The longer time points corresponded with a higher fluorescence because there was more time for the activated Cas12a to cleave the reporters.

Next, we normalized the inhibition curves to the corresponding positive and negative controls at each time point to facilitate comparisons across time points (Figure 2C). Assay incubation time did not have an impact on the normalized enzyme inhibition curves, which all overlapped with one another. We anticipate that the specific time point did not matter because Cas12a cleaves at a consistent rate and a longer assay duration simply results in more reporters being cleaved.^29^ Since all the time points displayed the same normalized fluorescence intensity curve, we chose 30 minutes for the final REACTR assay, as it provided a good tradeoff between maximizing the raw fluorescence signal and providing an assay time that is suitable for rapid POC testing. Our choice was also informed by a target product profile for an HIV medication adherence monitoring that recommends ≤ 60-minute analysis time.^15^

### DNA template length impacts REACTR sensitivity to AZT-TP

Next, we investigated the impact of DNA template length on REACTR’s ability to detect AZT-TP based on findings from our prior work with an intercalating dye reporting system.^20^ We designed DNA templates to vary the length of CTAA repeats between the RT primer binding region and the crRNA recognition region (Figure 3A, Supplementary Table 1). We chose CTAA as the repeating unit to minimize self-dimerization and secondary structure of the DNA template, which can inhibit or slow HIV RT activity. Enriching for adenosine bases also increases the likelihood of AZT-TP binding since the drug is a thymidine analog, thus providing more opportunities for AZT-TP incorporation and cDNA chain termination before activation of the crRNA-Cas12a complex.

**Figure 3.**
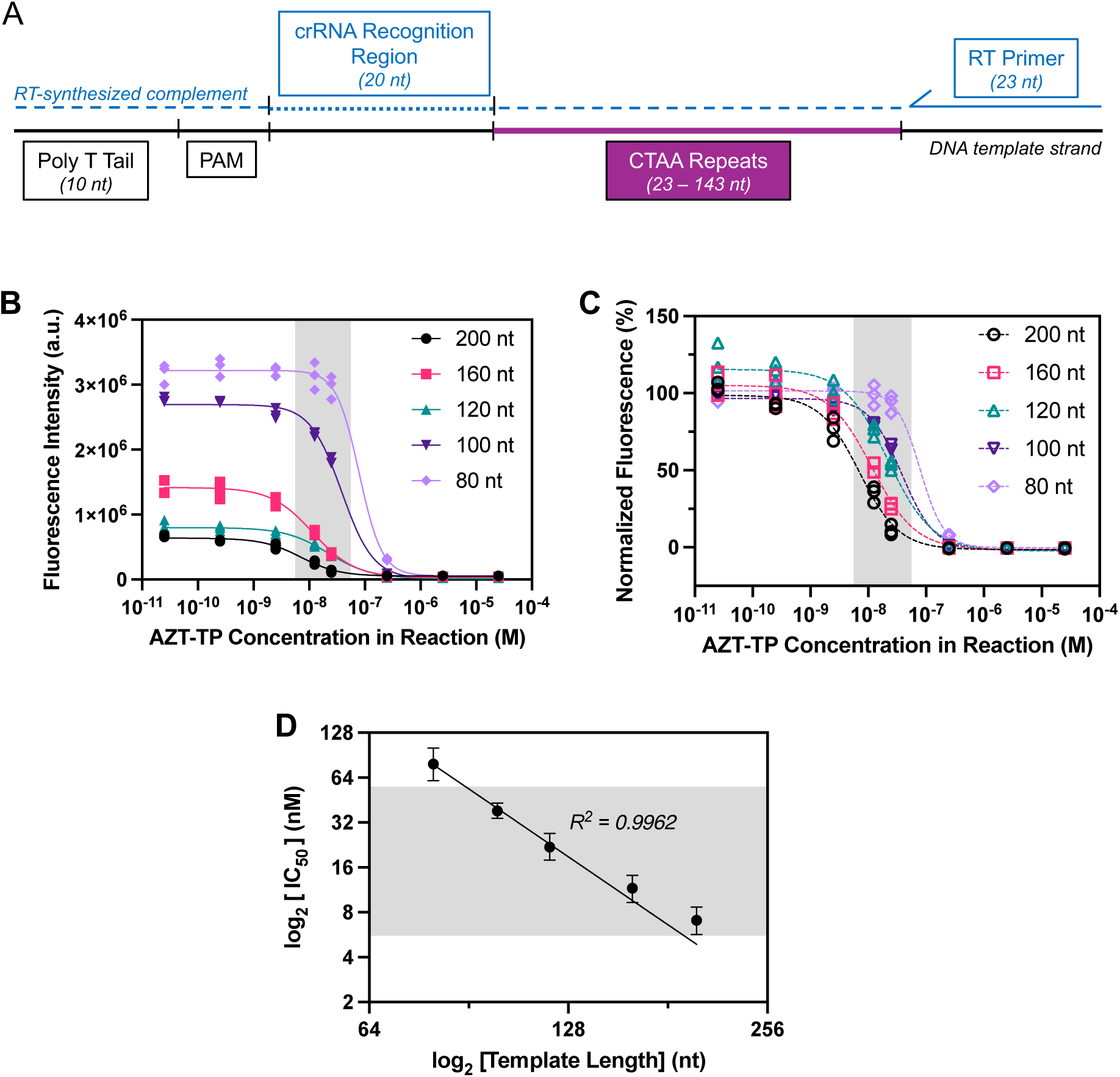
Impact of DNA Template Length on REACTR. **(A)** Diagram of the DNA template used for REACTR experiments. The template includes an RT primer binding region followed by CTAA repeats, ranging from 5 to 35 repeats, resulting in templates 80 to 200 nucleotides in length. The repeats are followed by the complement to the crRNA recognition sequence, a PAM terminating sequence, and a Poly T-tail. **(B)** Raw fluorescence data and **(C)** Normalized fluorescence data plotted for varying template lengths for AZT-TP concentrations (*N = 3)*. Data were analyzed at the 30-minute time point. **(D)** Calculated IC_50_ values as a function of DNA template length. Error bars indicate 95% confidence intervals. Grey bands indicate clinical range of AZT-TP that accumulates in white blood cells.^23^

We ran REACTR with DNA template lengths of 80, 100, 120, 160, and 200 nucleotides (nt) across a range of AZT-TP concentrations (Figure 3B). We saw a direct relationship between template length and amount of fluorescence generated in the 30 minutes of assay incubation. The shortest template, 80nt, had the highest fluorescence intensity compared to longer templates since it required the fewest nucleotide incorporation steps to reach the crRNA recognition region and activate Cas12a. Conversely, the 200nt template required the greatest number of nucleotide incorporation steps before the crRNA recognition region was fully synthesized and had the lowest observed fluorescence at the 30-minute timepoint. Four of the five templates tested followed this relationship between length and endpoint fluorescence, except for the 120nt template which had a slightly lower fluorescence than expected. Individual real-time fluorescence intensity for all the tested template lengths is provided in Supplemental Figure S2.

We normalized the inhibition curves to more easily compare effects of the DNA template length on REACTR’s sensitivity to AZT-TP in a clinical range of drug concentrations found in PBMCs.^20,23^ After normalization, we observed that as template length increased, inhibition curves shifted to lower AZT-TP concentrations (Figure 3C). This shift occurred because longer templates provided more binding sites for AZT-TP to be incorporated into the cDNA strand, increasing the likelihood of chain termination before reaching the crRNA recognition region. This is reflected in the 50% inhibition concentrations (IC_50_) values of the REACTR curves, which logarithmically decreased as template length increased (Figure 3D). These results agree with findings in our past work using REACT and intercalating dyes to measure NRTI drug levels.^20^ This ability to shift the REACTR curve allows us to design and optimize templates to measure clinically relevant concentrations of AZT-TP. We chose to use the 200nt template with the most binding regions to allow detection of the lowest concentrations of AZT-TP.

### REACTR retains sensitivity in presence of genomic DNA

AZT-TP accumulates in white blood cells; to release and detect AZT-TP, the cells must be lysed, which also releases genomic DNA (gDNA) found in the nucleus.^31,32^ Our previous work with enzyme inhibition assays for HIV drug level monitoring (REACT) used PicoGreen (PG), an intercalating dye, to enable fluorescent readout; however, PG indiscriminately binds to all double-stranded DNA, resulting in an inability to distinguish drug concentrations in the presence of gDNA (Supplemental Figure S3).^18^ REACTR selectively detects cDNA of interest using a CRISPR Cas reporter system, so we expect accurate results even in the presence of white blood cell lysate. We tested REACTR assay’s ability to distinguish between positive and negative controls with spiked gDNA (Figure 4A). We chose a biologically relevant concentration of gDNA in whole blood at 39.1 ng/µL^33^ and diluted the gDNA in nuclease-free water 1:4 and 1:10 to achieve final concentrations of 10 ng/µL and 4 ng/µL, respectively. The dilution factors were chosen to incorporate typical sample dilutions that have been used in our enzymatic assay.^18,19^ Figure 4A shows that the addition of gDNA had no impact on REACTR assay fluorescence, indicating that REACTR is highly selective for the DNA strand of interest even in a background of gDNA.

**Figure 4.**
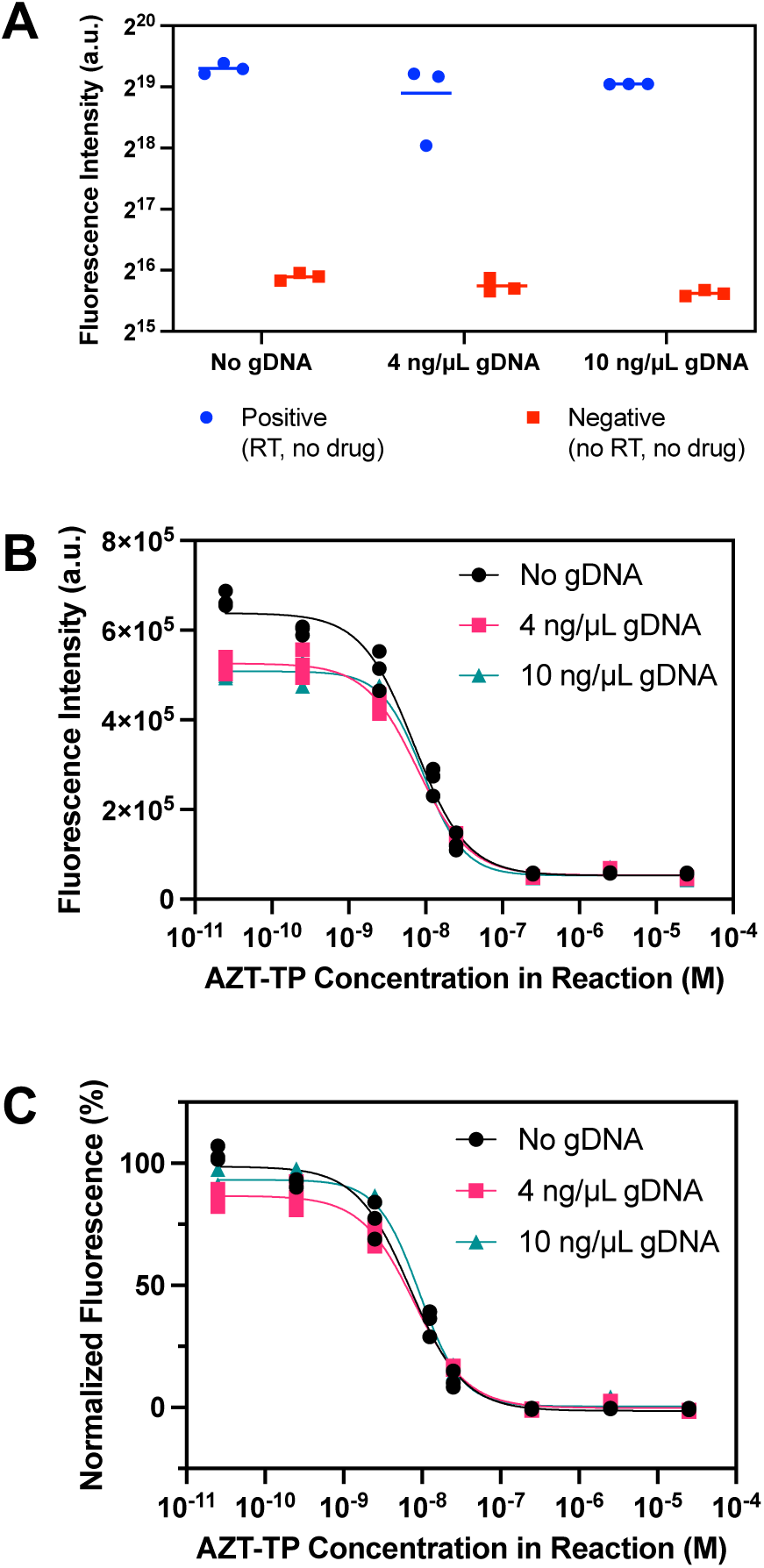
REACTR in the presence of genomic DNA. **(A)** Positive and negative controls at different genomic DNA (gDNA) concentrations. A solid line for each condition represents the mean (*N = 3)*. **(B)** Raw fluorescence data and **(C)** Normalized drug inhibition curves for AZT-TP concentrations for reactions containing no gDNA, 4 ng/µL of gDNA, and 10 ng/µL of gDNA (*N = 3)*.

Next, we tested REACTR assay’s sensitivity in differentiating various AZT-TP concentrations in the presence of gDNA. The raw drug inhibition curves showed slightly lower fluorescence in the presence of spiked gDNA (Figure 4B, Supplemental Figure S4). We hypothesize that this is because, unlike in the buffer condition, some of CRISPR collateral cleavage in the spiked gDNA condition cleaves gDNA instead and thus, does not result in fluorescence signal generation. After normalization, we observed no difference between the spiked gDNA and buffer reaction conditions, demonstrating that the REACTR assay can successfully detect AZT-TP with the same sensitivity at clinically relevant gDNA concentrations (Figure 4C).

### REACTR assay with PBMC lysate

Given that AZT-TP accumulates in PBMCs and must be lysed to access AZT-TP, we next investigated how the presence of PBMC lysate affected the REACTR assay. We lysed isolated PBMCs using the M-PER lysis buffer at a biologically relevant concentration (2E6 cells/mL)^34,35^ and selected dilution ratios (4%, 10%, 25%, and 100%) to represent different levels of sample preparation prior to REACTR. We confirmed that our standard lysis procedure released gDNA from PBMCs using PG intercalating dye (Supplemental Figure S5). We evaluated the impact of PBMC lysate on REACTR positive and negative controls (Figure 5A) and observed comparable fluorescence compared to the no lysate controls. As PBMC concentration increased, the fluorescence from negative controls (no RT, no drug) slightly increased, although they were still clearly differentiable from the positive controls (RT, no drug).

**Figure 5.**
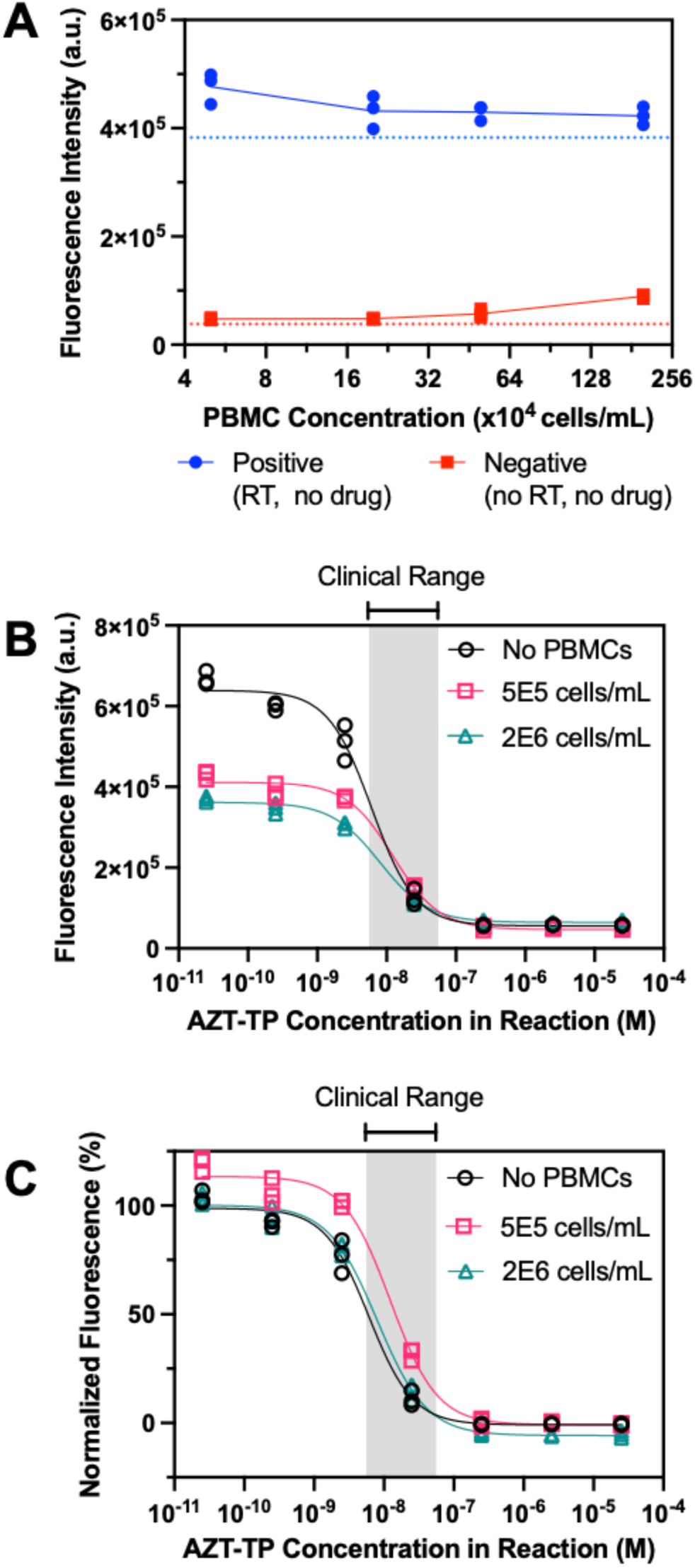
REACTR performance in PBMC lysate. **(A)** Positive and negative controls for different PBMC lysate dilutions ranging from 4% to 100% (*N = 3)*. The red and blue dotted lines represent the negative and positive buffer controls. **(B)** Fluorescence inhibition curves for varying AZT-TP concentrations with *N = 3.* Two PBMC lysate concentrations, 5E5 cells/mL and 2E6 cells/mL, were chosen and compared to a no PBMC lysate control. Data were analyzed 30 minutes after incubation. **(C)** Normalized drug inhibition curves. Grey bands indicate clinical range of AZT-TP that accumulates in white blood cells.^23^

We generated enzyme inhibition curves with the two highest PBMC concentrations tested (5E5 cells/mL and 2E6 cells/mL) to evaluate REACTR performance with the minimal amount of sample preparation (Figure 5B). We observed comparable gDNA concentrations between the gDNA and PBMC conditions, with there being 13.3ng/µL gDNA in 2E6 PBMCs and 3.3ng/µL gDNA in 5E5 PBMCs.^36^ Similar to REACTR results with spiked gDNA, raw fluorescence intensities with PBMC lysate were notably lower than in buffer (Figure 5B, Supplemental Figure S6). However, normalized inhibition curves were similar in PBMC lysate and in buffer (Figure 5C) and overlapped with the clinical range for AZT-TP.

We expect the enzyme inhibition curves to maintain the same sigmoidal slope even as PBMC lysate concentration increases. Sigmoidal enzyme inhibition curves are typically fit to four-parameter logistic regression curves with four key parameters: Hill slope, 50% inhibition concentration (IC50), top, and bottom. Of these parameters, the Hill slope which indicates the steepness of the sigmoidal curve and the IC50 which indicates the midpoint of the sigmoidal curve are the most informative, especially since the top and bottom parameters are typically normalized to 0% and 100%, respectively. When comparing the Hill Slope across the three conditions (buffer, 5E5 cells/mL, and 2E6 cells/mL) using a global least regression, we found no statistically significant difference (p=0.6892). The similar Hill slopes mean that our ability to distinguish low versus high drug levels on the enzyme inhibition curves remained similar regardless of PBMC lysate concentration. However, there was a statistically significant difference (p<0.0001) in the IC50 values across the 3 conditions. The IC50 values for buffer, 5E5 cells/mL, and 2E6 cells/mL were 6.1 nM (95% confidence interval [CI]: 4.9 – 7.7 nM), 12.3 nM (95% CI: 9.9 – 15.2 nM), and 8.0 nM (95% CI: 6.7 – 9.5 nM), respectively. Even though the 5E5 cells/mL condition had a higher IC50 than the other conditions, the midpoints of all the enzyme inhibition curves were within a ∼2X concentration range – which is much smaller than the expected differences in clinical drug levels. Taken together, our global least squares regression analyses across the 3 conditions suggest that the REACTR assay retains its capability to distinguish AZT-TP levels even in clinically relevant PBMC lysate conditions.

### Conclusions

We developed REACTR as a rapid enzymatic assay for real-time measurement of nucleotide analog drugs based on the detection of DNA synthesis by HIV RT enzyme using a CRISPR-Cas enabled readout. We characterized the effect of reaction time and DNA template length on REACTR assays using AZT-TP spiked in buffer. Unlike prior work that used intercalating dyes to measure HIV RT DNA synthesis, REACTR’s readout is specific to the synthesized DNA and is not compromised by background DNA present in clinical samples. REACTR selectively measured AZT-TP spiked into clinically relevant concentrations of genomic DNA and PBMC lysate. These results demonstrate the feasibility of using CRISPR-Cas-based system for measuring essential medications used to treat children living with HIV. The ability to monitor HIV drug levels quickly using readily available reagents will help to maintain therapeutic drug concentrations and minimize treatment failure, drug resistance, and medication side effects. Although the current iteration of REACTR requires fluorescence readout with a plate reader, future studies could use inexpensive portable readers^37^ or couple CRISPR reporters with instrument-free lateral flow readout.^38^

In addition, our proof-of-principle demonstration used pre-isolated PBMCS and did not include sample preparation from whole blood. Most CRISPR Cas readout demonstrations in the literature typically include sample preparation to isolate the analyte of interest and to minimize interference from sample matrices. The ideal sample preparation method would involve minimal purification/separation of the blood matrix components to reduce test complexity and cost. One recent report^39^ found that hemoglobin – a key component in red blood cell hemolysate – did not significantly diminish Cas 12 activity. We have also previously shown that blood dilution can minimize the non-specific inhibition of HIV RT by blood matrix components.^18,19^ We would first investigate dilution of whole blood and PBMC lysis using chemical methods^40^ as an initial simple sample preparation strategy. If additional sample preparation is required, we would explore alternate strategies like heating^24,41^ that have been shown to improve the accuracy and reproducibility of CRISPR readout without significantly increasing test complexity.

REACTR could significantly improve access to therapeutic drug monitoring in low-resource settings where HIV is endemic. REACTR can also be readily adapted to measure nucleotide analogs used in other HIV treatment regimens and other diseases (e.g. hepatitis^42^, cancer^43,44^) by designing DNA templates that account for the chemical structure of the drug of interest and the endogenous nucleotide that it mimics.^20^

## Methods

### REACTR Assay Workflow

The Cas12a-crRNA complex was created by combining 100 nM LbCas12a (M0653T, New England Biolabs) and 100 nM crRNA (Integrated DNA Technologies, see sequence in Table S1) in HIV RT buffer containing 50 mM Tris-HCl (77-86-1, Sigma-Aldrich) at pH 8, 50 mM Potassium Chloride (KCl) (60142-100ML-F, Sigma-Aldrich), 10 mM Magnesium Chloride (MgCl_2_) (7786-30-3, Sigma Aldrich), 2 mM dithiothreitol (DTT) (3483-12-3, Sigma-Aldrich), and 0.06% Triton X-100 (9036-19-5, Sigma-Aldrich). The reagents were incubated for 15 minutes at 25°C.

AZT-TP (B1331-007254, BOC Sciences) was diluted in nuclease-free water at varying concentrations from 10^-4^ to 10^-10^ M. 40 µL REACT assays containing 10 µL of the drug dilution, 10 µL of the Cas12-crRNA complex, 2 µL (20.2 U/µL) of HIV RT enzyme (LS005009, Worthington Biochemical), and 18 µL of reaction master mix were added to black, flat-bottom polystyrene 96-well plate with nonbinding surfaces (3650, Corning). The master mix consisted of 0.2 µL (1 M) MgCl_2_ and 0.2 µL (1 M) DTT, 0.2 µL (100 µM) deoxynucleotides (dNTP) (N0447L, New England Biolabs), 0.1 µL (20 µM) RT primer, 0.1 µL (2 µM) DNA template, and 1 µL (20 µM) reporters (Integrated DNA Technologies), 4 µL RT buffer, and 12.2 µL nuclease-free water. HIV RT enzyme was added last to initiate the reaction. Assays with drug dilutions were compared against “no-drug” positive controls and “no-enzyme” negative controls.

The positive control replaced the drug dilutions with nuclease-free water while the negative controls replaced the drug dilutions with nuclease-free water and replaced HIV RT with RT buffer. Reactions were incubated at 37 °C in a microplate reader (SpectraMax iD3, Molecular Devices) for 90 minutes. Fluorescence measurements were taken every two minutes during the incubation period, with one ten-second shake before the first fluorescence measurement.

### Template Design

DNA templates and primer, which were purchased from Integrated DNA Technologies (IDT), were designed to minimize secondary structures and self-dimerization. The DNA templates were designed to have a primer binding region followed by CTAA repeats, the complement crRNA recognition region, the PAM sequence required for Cas12a cleavage, and a 10nt Poly-T tail. Prior work showed that addition of a poly-T tail near the activation sequence increased fluorescence readout from the CRISPR Cas complex.^28^ The CTAA repeating region ranged from 5 to 35 repeats to reach 80nt and 200nt, respectively. The CTAA repeat was designed to include more adenosine bases because AZT-TP is a deoxythymidine triphosphate (dTTP) analog and will bind to A’s in the template. The final designs of the templates and primer are listed in Table S1.

### REACTR with spiked genomic DNA

100 µg Human Mixed Genomic DNA (G3041, Promega) was purchased and diluted in nuclease-free water prior to experiments. Genomic DNA was diluted to final concentrations of 4 ng/µL and 10 ng/µL in the master mix. REACTR was completed as described above, except genomic DNA was added to the master mix as an extra component that replaced some of the nuclease-free water.

### REACTR with spiked PBMC lysate

1E7 cells/mL human PBMCs (HUMANHLPB-0001146, BioIVT) were resuspended in 1 mL of M-PER Mammalian Protein Extraction Reagent (78501, Thermo Fisher Scientific) for lysis. The lysed PBMCs were shaken at 250 rotations per minute (rpm) for 10 minutes and were diluted with TE Buffer to an equivalent concentration of 2E6 cells/mL, the estimated concentration of PBMCs in 1 mL of whole blood.^34^ PBMC lysate was then diluted with TE buffer at various (volume/volume) dilution ratios 4%, 10%, 25%, and 100% to represent different degrees of sample preparation prior to testing. We then added 10 µL of AZT-TP spiked in the PBMC lysate to the reaction mixture so the final PBMC lysate concentration was 1/4^th^ the concentration in the assay. We prepared serial dilutions of clinically relevant levels of AZT-TP with concentrations ranging from 1E-4 to 1E-10 M. The 200nt DNA template was chosen as the template for the master mix, while the rest of the components remained the same. The RT and the Cas complex were prepared as described above and reactions were incubated also as described above.

### Data analysis

The enzyme inhibition curves were generated using REACTR fluorescence at the 30-minute time point. The fluorescence intensities were then normalized using the average of the positive and negative controls as maximum and minimum values, respectively.^45^ The normalized curves were fitted to four-parameter logistic regression curves using GraphPad Prism 10 (GraphPad Software). To determine whether the IC50 and Hill slope coefficients differed significantly between PBMC lysate concentrations, we applied the extra sum-of-squares F-test using GraphPad Prism10.

## Declarations

### Conflicts of Interest

M.A.S., M.M.C., Q.W., C.R., B.R.L., and A.O.O. are inventors on a patent filed (63/723495) based on this work. A.O.O. is an inventor on a patent filed (PCT/US2020/037609) based on related work on enzymatic assays for measuring reverse transcriptase inhibitors. B.R.L. holds equity in a startup company that has licensed related technology and supports ongoing work in the B.R.L. laboratory at the University of Washington. B.R.L serves as a scientific advisor for the company. The company played no role in the funding, study design, data analyses, or reporting of results for this study.

### Author Contributions

**Maya Singh:** Conceptualization; Formal Analysis; Investigation; Methodology; Writing – Original Draft Preparation; Writing – Reviewing and Editing; Visualization; Project Administration. **Megan Chang:** Conceptualization; Funding Acquisition; Methodology; Writing – Reviewing and Editing; Visualization. **Qin Wang** Methodology; Resources. **Catherine Rodgers:** Methodology. **Barry Lutz:** Resources.

**Ayokunle Olanrewaju:** Conceptualization; Funding Acquisition; Methodology; Project Administration; Resources; Writing – Reviewing and Editing.

## Acknowledgements

The authors are grateful for funding from the Washington Research Foundation. We thank the Washington Entrepreneurial Research Evaluation and Commercialization Hub (WE-REACH), the National Heart, Lung, and Blood Institute (NHLBI) {NIH Grant # U01 HL152401}, and matching grant partners for supporting A.O.O, M.A.S, and M.M.C. The content of this research is solely the responsibility of the authors and does not necessarily represent the official views of the National Institutes of Health. This work was supported by NIH/NIAID grants R33AI140460. We thank Cosette Craig and Cara Brainerd for their feedback on experiments and the manuscript.

