## Supplemental Information for "Rapid enzymatic assay for antiretroviral drug monitoring using CRISPR-Cas12a enabled readout"

### Table of Contents

| **Supplementary Information Summary** **………………………………………………………………** | **S2** |
| --- | --- |
| **Table S1.** DNA and RNA Sequences for Template Design**…………………………………………...** | **S3** |
| **Figure S1.** REACTR Real-time Performance Curve with Error Bars**………………………………..** | **S4** |
| **Figure S2.** REACTR Real-time Performance for each CTAA Template**………………………….....** | **S5** |
| **Figure S3.** Addition of Genomic DNA to Intercalating Dye Based REACTR**……………………….** | **S6** |
| **Figure S4.** REACTR Real-time Performance Curves for Varying Genomic DNA Concentrations**……………………………………………………………………………………………..** | **S7** |
| **Figure S5.** Varying PBMC Lysate Concentrations with PicoGreen Readout**……………………….** | **S8** |
| **Figure S6.** REACTR Real-time Performance Curves for Varying PBMC Lysate Concentrations**……………………………………………………………………………………………..** | **S9** |

**Supplementary Information**

Supplementary information includes the following: DNA and RNA sequences used during the experimental workflow, REACTR real-time curves with standard error bars, REACTR real-time curves for varying CTAA template lengths, a direct comparison between REACTR and an intercalating dye-based readout, REACTR real-time curves for varying genomic DNA concentrations spiked into the assay, varying PBMC lysate concentrations with a PicoGreen readout, and REACTR real-time curves for varying PBMC lysate concentrations.

**Table S1. DNA and RNA Sequences for Template Design.**

| ***SEQUENCES*** | |
| --- | --- |
| **Names** | **Sequence** |
| 80 nt CTAA Template | TTTTTTTTTTTTTGATGATGTGAAGGTGTTGTCGCTAACTAACTAACTAACTAACTAACTAACTAACTAACTAACTACTATCTTTCCTCTTAATTCGACG |
| 100 nt CTAA Template | TTTTTTTTTTTTTGATGATGTGAAGGTGTTGTCGCTAACTAACTAACTAACTAACTAACTAACTAACTAACTAACTAACTAACTAACTAACTAACTACTATCTTTCCTCTTAATTCGACG |
| 120 nt CTAA Template | TTTTTTTTTTTTTGATGATGTGAAGGTGTTGTCGCTAACTAACTAACTAACTAACTAACTAACTAACTAACTAACTAACTAACTAACTAACTAACTAACTAACTAACTAACTAACTACTATCTTTCCTCTTAATTCGACG |
| 160 nt CTAA Template | TTTTTTTTTTTTTGATGATGTGAAGGTGTTGTCGCTAACTAACTAACTAACTAACTAACTAACTAACTAACTAACTAACTAACTAACTAACTAACTAACTAACTAACTAACTAACTAACTAACTAACTAACTAACTACTATCTTTCCTCTTAATTCGACG |
| 200 nt CTAA Template | TTTTTTTTTTTTTGATGATGTGAAGGTGTTGTCGCTAACTAACTAACTAACTAACTAACTAACTAACTAACTAACTAACTAACTAACTAACTAACTAACTAACTAACTAACTAACTAACTAACTAACTAACTAACTAACTAACTAACTAACTAACTAACTAACTAACTAACTAACTACTATCTTTCCTCTTAATTCGACG |
| DNA Primer | CGTCGAATTAAGAGGAAAGATAG |
| crRNA Recognition Region | TACTACACTTCCACAACAGC |
| Fluorescence Reporter | /56-FAM/TTATT/3IABkFQ/ |
| crRNA | rUrArA rUrUrU rCrUrA rCrUrA rArGrU rGrUrA rGrArU rArUrG rArUrG rUrGrA rArGrG rUrGrU rUrGrU rCrG |

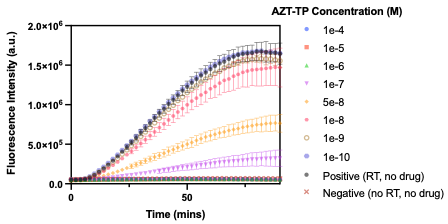

**Figure S1. REACTR Real-time Performance Curve with Error Bars.** Real-time REACTR curves with varying concentrations of AZT-TP. Each data point represents the mean fluorescence measurements with error bars representing standard deviation (*N = 3*)*.*

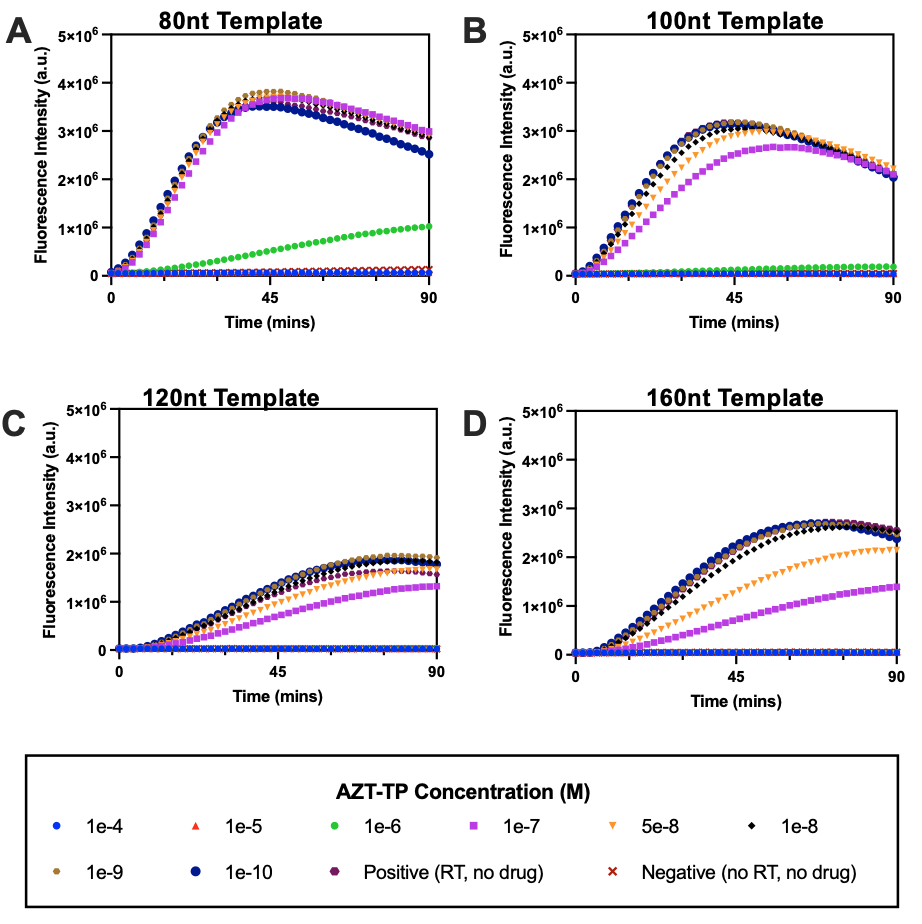

**Figure S2. REACTR Real-time Performance for each CTAA Template.** **(A)** Real-time REACTR curves with varying concentrations of AZT-TP. Each data point represents the mean fluorescence measurements (*N = 3*) for 80 nt template, **(B)** 100 nt template, **(C)** 120nt template, and **(D)** 160nt template.

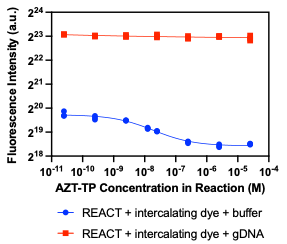

**Figure S3. Addition of Genomic DNA to Intercalating Dye Based REACTR.** We used Picogreen as the intercalating dye to measure the fluorescence intensity after a 30-minute incubation period for the 200nt CTAA template. The addition of gDNA caused an increase in fluorescence and had indistinguishable signal between AZT-TP concentrations. We saw distinguishable fluorescence values for each AZT-TP concentration in the Picogreen only condition.

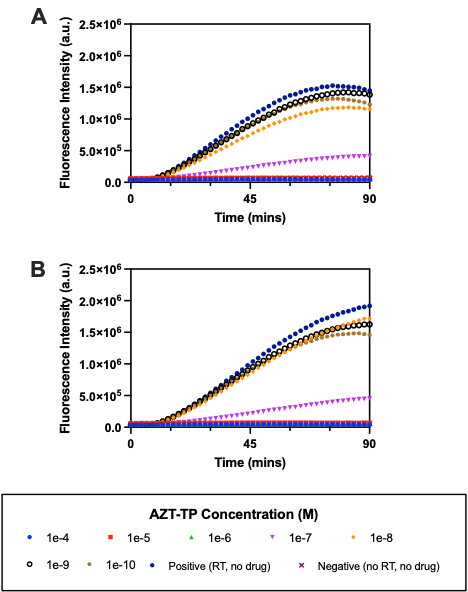

**Figure S4. REACTR Real-time Performance Curves for Varying Genomic DNA Concentrations.** **(A)** Real-time REACTR curves spiked with 4 ng/µL genomic DNA with varying concentrations of AZT-TP. Each data point represents the mean fluorescence measurements (*N = 3*)*.* **(B)** Real-time REACTR curves spiked with 10 ng/µL genomic DNA with varying concentrations of AZT-TP. Each data point represents the mean fluorescence measurements (*N = 3*)*.*

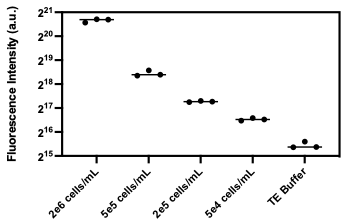

**Figure S5. Varying PBMC Lysate Concentrations with PicoGreen Readout.** Concentrations of PBMC lysate starting from 2E6 cells/mL were diluted with TE Buffer. Since PBMC lysate has double stranded DNA, we expected to see a decrease in fluorescence as we decreased the PBMC lysate concentration. We compared the fluorescence intensity with a TE buffer only condition. PBMC lysate was incubated with PicoGreen at 37^o^C for one minute before fluorescence measurements were taken.

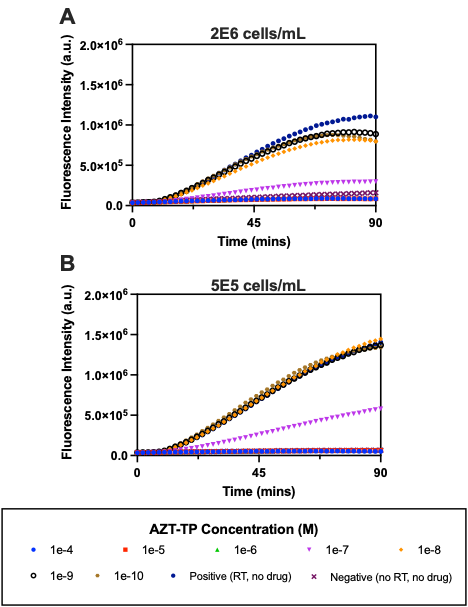

**Figure S6. REACTR Real-time Performance Curves for Varying PBMC Lysate Concentrations.** **(A)** Real-time REACTR curves spiked with 2E6 cells/mL PBMC lysate with varying concentrations of AZT-TP. **(B)** Real-time REACTR curves spiked with 5E5 cells/mL PBMC lysate genomic DNA with varying concentrations of AZT-TP. Each data point represents the mean fluorescence measurements (*N = 3*)*.*
